# An ensemble of interconverting conformations of the elemental paused transcription complex creates regulatory options

**DOI:** 10.1101/2022.09.11.507475

**Authors:** Jin Young Kang, Tatiana V. Mishanina, Yu Bao, James Chen, Eliza Llewellyn, James Liu, Seth A. Darst, Robert Landick

## Abstract

Transcriptional pausing underpins regulation of cellular RNA synthesis but its mechanism remains incompletely understood. Sequence-specific interactions of DNA and RNA with the dynamic, multidomain RNA polymerase (RNAP) trigger reversible conformational changes at pause sites that temporarily interrupt the nucleotide addition cycle. These interactions initially rearrange the elongation complex (EC) into an elemental paused EC (ePEC). ePECs can form longer-lived PECs by further rearrangements or interactions of diffusible regulators. For both bacterial and mammalian RNAPs, a half-translocated state in which the next DNA template base fails to load into the active site appears central to the ePEC. Some RNAPs also swivel interconnected modules that may stabilize the ePEC. However, it is unclear if swiveling and half-translocation are requisite features of a single ePEC state or if multiple ePEC states exist. Here we use cryo-EM analysis of ePECs with different RNA–DNA sequences combined with biochemical probes of ePEC structure to define an interconverting ensemble of ePEC states. ePECs occupy either pre- or half-translocated states but do not always swivel, indicating that difficulty in forming the post-translocated state at certain RNA–DNA sequences may be the essence of the ePEC. The existence of multiple ePEC conformations has broad implications for transcriptional regulation.

**SIGNIFICANCE:** Transcriptional pausing provides a hub for gene regulation. Pausing provides a timing mechanism to coordinate regulatory interactions, co-transcriptional RNA folding and protein synthesis, and stop signals for transcriptional termination. Cellular RNA polymerases (RNAPs) are complex, with multiple mobile modules shifting positions to control its catalytic activity and pause RNAP in response to DNA-encoded pause signals. Understanding how these modules move to enable pausing is crucial for a mechanistic understanding of gene regulation. Our results clarify the picture significantly by defining multiple states among which paused RNAP partitions in response to different pause signals. This work contributes to an emerging theme wherein multiple interconverting states of the RNAP proceed through a pathway (e.g., initiation or pausing), providing multiple opportunities for regulation.

## INTRODUCTION

Transcriptional pausing is an evolved feature of all cellular DNA-dependent RNA polymerases (RNAPs) responsible for the regulated expression of genes. At pause sites, interactions of RNAP with certain RNA–DNA sequences temporarily halt transcription for tens to thousands of times longer than the 10-20 ms average nucleotide addition cycle (NAC) (for *E. coli* RNAP). These delays allow time for folding of nascent RNA into biologically active forms, interactions with diffusible regulators, coupling with translation or splicing, termination of transcription, or other events that regulate gene expression (1, 2).

A now widely accepted model posits that RNA–DNA interactions put RNAP into an initially paused off-pathway state called the elemental pause (1, 3–5). Formation of the elemental paused elongation complex (ePEC) competes with the NAC rather than halting all RNAPs (Fig. 1A). Conformational fluctuations in the complex, multidomain RNAP, modulated by RNA–DNA sequence, govern this competition and ePEC lifetime. The ePEC can rearrange into longer-lived pauses by backtracking of the RNA and DNA chains, by interactions with nascent RNA secondary structures (pause hairpins; PHs) (6, 7), or by interactions with diffusible regulators (e.g., NusA, ppGpp, or pro-pausing NusG/Spt5) (8–10). RNA structures and regulators also can shorten or suppress elemental pausing (e.g., HK022 *put* RNA and λQ antiterminators or anti-pausing NusG/Spt5) (11–14). The elemental pause model arose from the observation of residual pausing when the *E. coli his* operon leader PH was deleted and from detection of non-backtracked paused states in single-molecule experiments (3, 15, 16). The kinetic characteristics of elemental pausing are confirmed by multiple studies (17–21). Genome-scale analyses reveal a consensus sequence for strong elemental pauses recognized by diverse RNAPs, including during initial transcription prior to promoter escape and for mammalian RNAPII (5’-GGnnnnnntg**YR**ccc, where YR corresponds to the pause RNA 3’ end and incoming NTP) (22–26).

**FIGURE 1.**
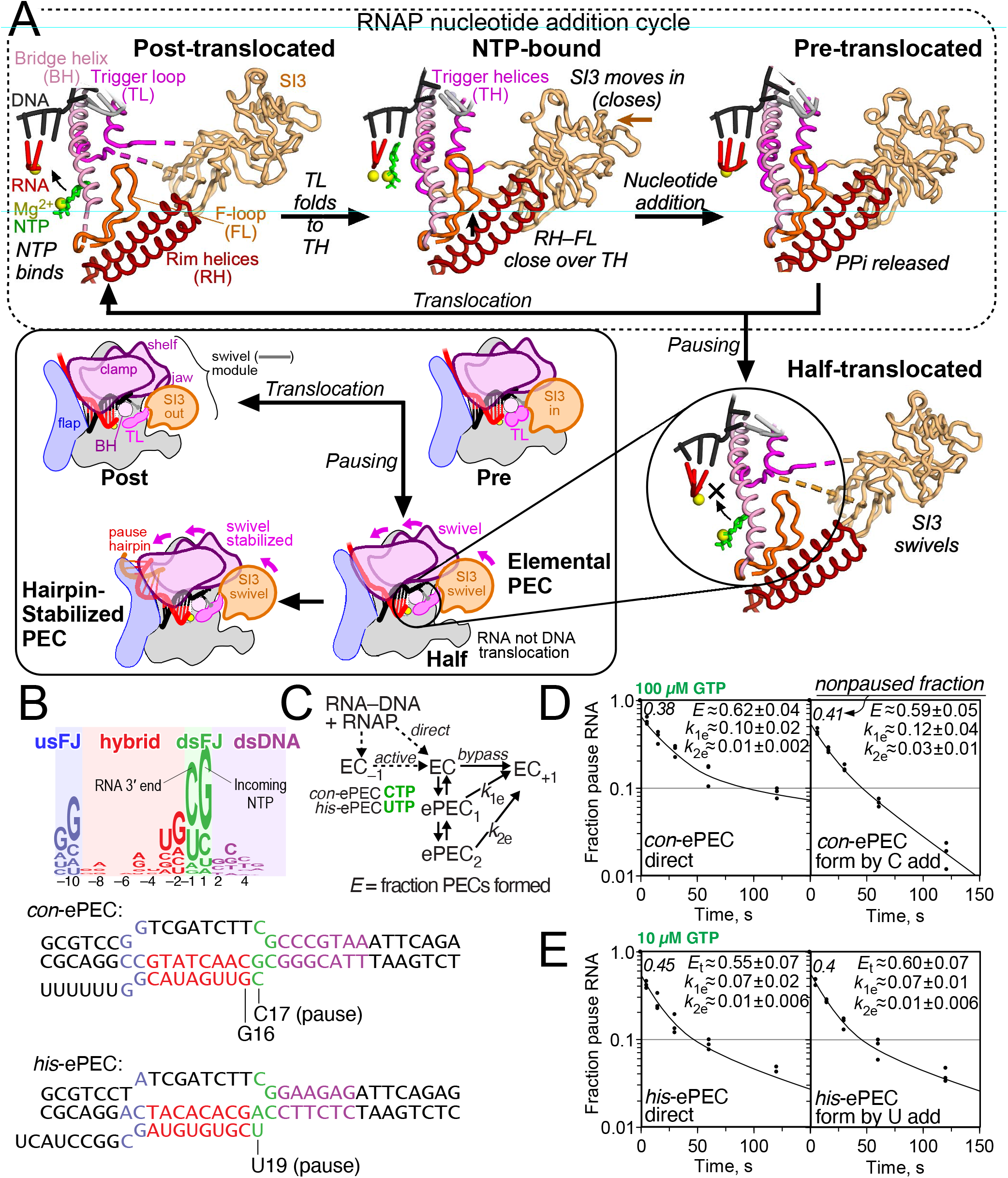
Elemental pausing and the RNAP nucleotide addition cycle. (**A**) Active site changes during the RNAP nucleotide addition cycle. Both the bridge helix (BH) and trigger loop (TL) undergo conformational changes upon NTP binding, catalysis, PPi release, and translocation. The TL folds from a flexible loop to a helical hairpin that, in *E. coli* and related bacteria, contains a large (188aa) insertion called sequence insertion 3 (SI3). The rim helices (RH) and F-loop (FL) shift positions when the TL folds. Pause sequences induce formation of an elemental pause conformation in which the RNA but not DNA are translocated (half-translocated). The ePECs can further rearrange by swiveling of a swivel module that inhibits TL folding via steric effects on SI3 and nascent RNA hairpin formation that stabilizes the swiveled conformation. The structural models are based on PDB 6ALF (post-translocated), 6RH3 (NTP-bound), con-ePEC-fTL (pre-translocated; this work), and *his*-ePEC-ufTL (half-translocated; this work)(9, 27). (**B**) A consensus elemental pause sequence has been defined by NET-seq in *E. coli*(22). The consensus consists of four distinct elements, the upstream fork junction (usFJ), RNA–RNA hybrid, downstream fork junction (dsFJ), and downstream duplex DNA. Two ePECs are relatively well studied, the consensus ePEC (*con*-ePEC) and the ePEC formed prior to pause hairpin formation in the *his* operon leader region (*his*-ePEC). Sequences shown are the scaffolds used for cryo-EM analysis in this work. (**C**) Two methods to assemble ePECs were used in this work and compared kinetically. Active ePEC formation was accomplished reaction of CTP or UTP to ECs assembled one nt upstream from the pause sites. Direct ePEC formation was accomplished by mixing RNAP with an RNA–t-DNA scaffold followed by binding of nt-DNA without any nucleotide addition. (**D**) Kinetic comparison of reaction with 100 μM GTP of *con*-ePECs formed by the direct or active methods. Both methods yielded biphasic pause escape rates. Equivalent fractions of slow and more slowly escaping ePECs formed by each method, but the escape rates were slight faster for the active ePECs. (**D**) Kinetic comparison of reaction with 10 μM GTP of *his*-ePEC formed by the direct or active methods. Both methods yielded biphasic pause escape rates and equivalent fractions and escape rates for the slow and more slowly escaping ePECs.

Cryo-EM of both *E. coli* and mammalian PECs reveal a half-translocated RNA–DNA hybrid (Fig. 1A) (9, 27, 28). In the half-translocated state, the recently added 3’ RNA nt has cleared the active site to allow binding of the next NTP substrate. However, the template DNA (t-DNA) base needed to specify the next NTP remains paired to the non-template DNA (nt-DNA) strand in the downstream DNA channel. Completion of DNA translocation to allow NTP binding appears to limit escape from the ePEC and reentry into the NAC (19).

In the paused state, modest rotation of a swivel module of RNAP (by ~1.5°–6° relative to an NTP-bound EC) (29) appears to inhibit RNAP motions required for completion of translocation, NTP binding, and catalysis (7, 9, 30). The swivel module comprises ~33% of the mass of bacterial RNAP and consists of the clamp, shelf, dock, jaw, β’C-term, and SI3 in *E. coli* RNAP (see Table S1 for RNAP structural modules). Weak swiveling (~1.5° average) is evident in a low-resolution (~5.5 Å) ePEC structure and detectable even in canonical ECs not bound to transcription factors or NTP (9, 30). Pronounced swiveling (~4.5–6°) occurs when a PH forms in RNAP’s RNA exit channel, NusA is bound, or both. For *E. coli* RNAP, which contains the 188-aa SI3 insertion in the trigger loop (TL) (31), pronounced swiveling appears incompatible with SI3 movements required for TL folding and bridge helix (BH), rim helices (RH), and F-loop (FL) movements that position NTP for nucleotide addition (Fig. 1A) (7, 29, 32).

Results to date suggest half-translocation and swiveling are characteristic of the paused state, but it remains unclear if all PEC states are half-translocated and swiveled, if multiple PEC states interconvert, and if different pause signals may generate different pause states. To gain greater insight into elemental pause state(s), we used cryo-EM and biochemical probes of RNAP conformation and translocation to examine ePECs formed on a strong, consensus elemental pause signal (*con*-ePEC) and a weaker signal that forms ePECs prior to stabilization by a PH in the *his* biosynthetic operon leader region (*his*-ePEC). We find that pause sequences lead to both pre- or half-translocated states, the common feature being inhibition of achieving the post-translocated state. In addition, different ePECs show differences in the global conformation of RNAP. Multiple states are observed on each pause sequence, and our kinetic modeling suggests how these various states might interconvert upon RNAP entry into a pause.

## RESULTS

### Multiple interconverting pause states form *in vitro*

To provide a kinetic framework for structural analysis of elemental pause states, we first compared formation and escape kinetics of the long-lived *con*-ePEC (19, 22) and the shorter-lived *his*-ePEC lacking the PH (7, 9, 15) (Fig. 1B; see Table S2 for oligonucleotides and plasmids used in this study). We examined both ePECs using two different ways to make them (Fig. 1C). One method, often used for cryo-EM, was to form ePECs by direct reconstitution from RNAP mixed with RNA and DNA strands (Fig. 1C; direct). The other method was to form the same ePECs by nucleotide addition from the ECs reconstituted at the –1 position by incubation with CTP (*con*-ePEC) or UTP (*his*-PEC) (Fig. 1C; active). Consistent with prior analyses (15, 19), both *con*-ePEC and *his*-ePEC exhibited biphasic apparent rates of pause escape and a non-paused subpopulation for both types of complexes (Figs. 1D,E). Biphasic escape rates reflect formation of multiple ePEC states. The non-paused subpopulations reflect fractions of ECs that do not pause and confirm that ePECs form as off-pathway states. Also consistent with prior analyses, *con*-ePEC paused states were roughly ten times longer lived than *his*-ePEC states. This conclusion derives from *con*-ePEC at 100 μM GTP and *his*-ePEC at 10 μM GTP exhibiting similar pause escape profiles (Figs. 1D,E) and assuming half-maximal [NTP]s are mM (22). Slight differences with prior analyses and between the two types of complexes for *con*-ePEC reflect the known kinetic heterogeneity of ePECs. Different RNAP preps and even different GTP concentrations can shift the distribution of ePECs among paused (and non-paused) states (19). The structural difference between fast and slow escaping ePECs has been unclear to date but is unlikely to involve backtracking. A 1-base pair (bp) backtrack is possible on the *con*-ePEC scaffold but it does not contribute to slow escape rates (19). Even 1-bp backtracking is strongly disfavored on the *his*-ePEC scaffold by an RNA–DNA base mismatch at –11 (Fig. 1B). We concluded that ePECs formed either by direct reconstitution or by one round of nucleotide addition were kinetically similar and were both suitable for structural analyses by cryo-EM.

### The *con*-ePEC occupies pre-translocated states that differ in TL folding and SI3 location

Understanding of PEC structure to date has relied on comparison of directly reconstituted PECs to non-paused ECs formed on unrelated DNA–RNA sequences. To compare the *con*-ePEC to a mechanistically related EC, we formed *con*-ePECs by single-round nucleotide addition at 23 °C with 200 μM CTP from 10 μM *con*-ePEC_−1_ (i.e., actively formed *con*-ePECs from a reconstituted EC poised 1 bp preceding the pause). *con*-ePEC formed rapidly. Even after 14 s and unchanged for up to 2 min when incubated with 10 μM GTP at 23 °C, *con*-ePEC exhibited indistinguishable biphasic escape kinetics with ~8% non-paused ECs (Fig. S1). Thus, at least two kinetically distinguishable *con*-ePEC states formed quickly are remained unchanged in proportion for at least 2 min.

Guided by these results, we determined cryo-EM structures for both *con*-ePEC_−1_ and *con*-ePEC using >300K polished particles for each. *con*-ePEC was formed by adding 200 μM CTP to 10 μM *con*-ePEC_−1_ and plunge freezing in liquid ethane after 14s at 23°C (Fig. 2). In each case, we used 3D classification to identify complexes containing intact scaffold (Fig. S2; Table S3). As expected, *con*-ePEC_−1_ was post-translocated with an unfolded TL (Fig. 2A). It generally resembled the conformation of a previously determined post-translocated EC structure containing an RNA 3’ A in the *i* site and t-DNA G in the open *i+1* site (27) (vs. 3’ rG and t-DNA C for *con*-ePEC_−1_).

**FIGURE 2.**
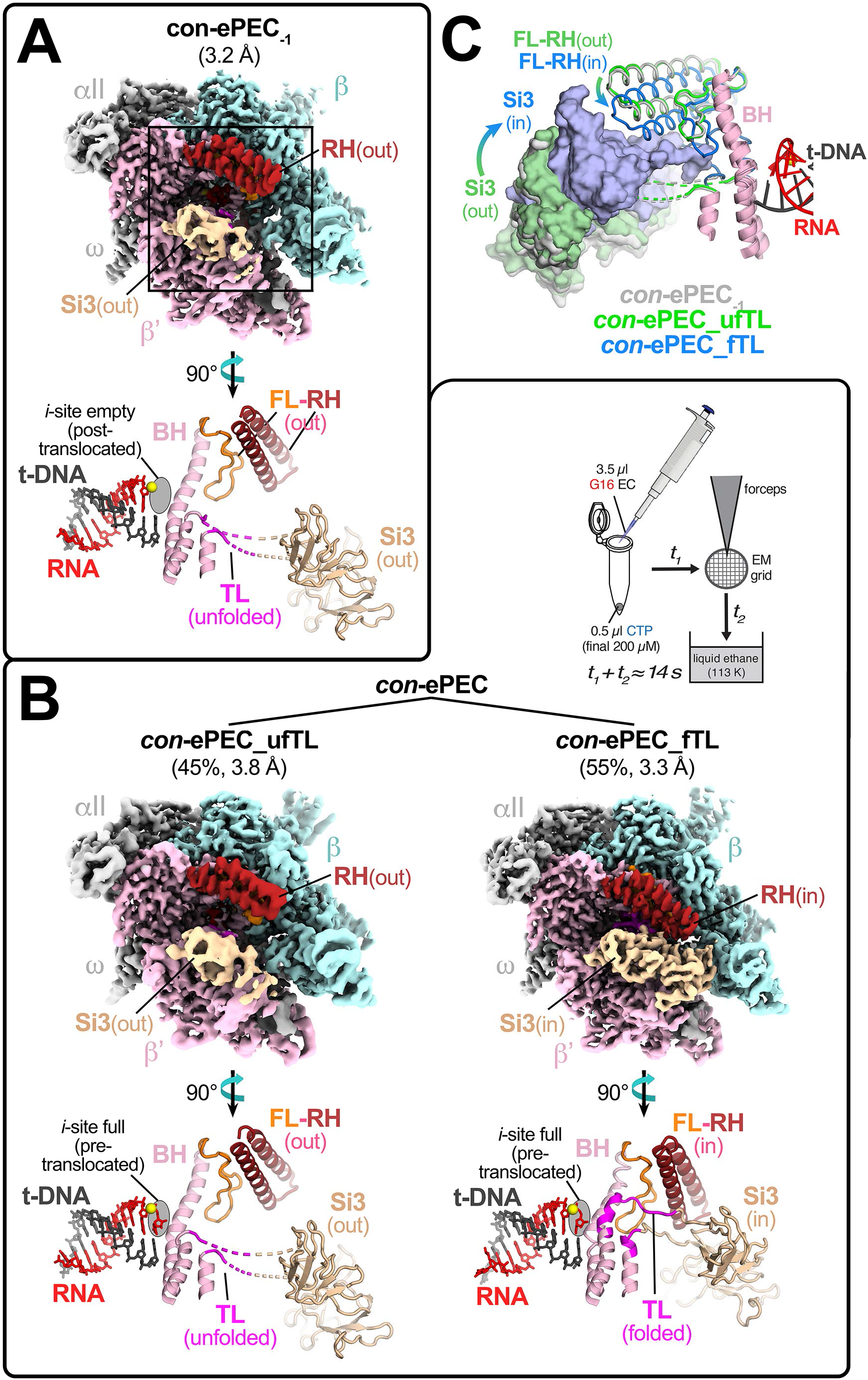
Cryo-EM analysis of actively formed *con*-ePEC. (**A**) The overall cryo-EM structure and active-site conformation of *con*-ePEC_−1_. The cryo-EM map is colored by RNAP subunit or feature with β’ in light pink, β in pale cyan, a and ω in light gray, SI3 in wheat, and the rim helices in dark red. An enlarged and rotated region of *con*-ePEC_−1_ around the active site is shown using secondary structure cartoons below the cryo-EM map with additional features F-loop in orange, bridge helix (BH) in light pink, trigger loop (TL) in light pink or magenta (flexible region), RNA (red), t-DNA (gray), and active-site Mg^2+^ (yellow sphere). (**B**) Inset (upper right of panel): *con*-ePECs were formed by ~14 s reaction at ~23 °C of directly reconstituted *con*-ePEC_−1_ with 200 μM CTP that included time on cryo-EM grid before plunge freezing in liquid ethane. The overall cryo-EM structures and active-site conformations of *con*-ePEC_ufTL and *con*-ePEC_fTL are shown, with cryoEM maps colored by RNAP subunits and features as described in panel (A). The percentages refer to the relative amounts of the two structures observed by cryo-EM (Figure S3). Enlarged and rotated regions around the active site are shown for each *con*-ePEC structure below the cryo-EM maps as described for *con*-ePEC_−1_ in panel (B). (**C**) Comparison of the locations of SI3, RH, and FL in *con*-ePEC_−1_ (light gray), *con*-ePEC_ufTL (green), and *con*_ePEC-fTL (light blue). The BH and TL-helices (light pink) are shown for reference.

*con*-ePEC was exclusively pre-translocated, in contrast to the half-translocated *his*-ePEC observed previously (7, 9). The pre-translocated *con*-ePECs sorted into two distinct classes. One subpopulation (*con*-ePEC_ufTL, ~45% of particles) contained an unfolded TL and SI3 in the open position (Fig. 2B). A second subpopulation (*con*-ePEC_fTL, ~55% of particles) contained a folded TL with SI3 in the closed position (similar to a previously described pre-translocated initiation complex except lacking σ^70^) (32, 33). In *con*-ePEC_fTL, the RH-FL module was rotated 11.9° towards the top of the folded TL (Fig. 2C). Thus, *con*-ePEC_fTL resembled most closely an NTP-bound EC in which the TL was folded and the RH-FL module was rotated (PDB 6RH3) (29). However, *con*-ePEC_fTL and *con*_ePEC_ufTL differed from each other at the upstream fork junction (usFJ). In *con*-ePEC_fTL, the RNA–DNA hybrid was 10 bp; the −11 RNA and partner t-DNA nucleotides were separated. In *con*-ePEC_fTL, the hybrid was nearly 11 bp because the −11 RNA and t-DNA nucleotides rotated back toward the hybrid on the main cleft side of the lid, approaching bp distance (~6 Å) (Fig. S3). Assuming *con*-ePEC_fTL is the first paused state formed after nucleotide addition, formation of a partial −11 bp in *con*-ePEC_ufTL may reflect failure to translocate upon TL unfolding and a conformational shift that, instead, allows the usFJ to rearrange. Formation of this −11 near-bp may stabilize the pretranslocated *con*-ePEC and explain the conservation of the strong −11 rG–dC bp in the consensus ePEC pause signal (22, 24).

These *con*-ePEC structures represent new paused conformations in which the EC remains pre-translocated with multiple TL conformations. Occupancy of predominantly pre-translocated rather than half-translocated states by the *con*-ePEC is consistent with prior findings that it is sensitive to pyrophosphorolysis and that the incoming t-DNA base is still paired to the nt-strand (19, 22).

### The *his*-ePEC is mostly half-translocated and swiveled but includes novel pre-translocated states

Although the differences between the *con*-ePEC and previously determined *his*-ePEC are striking, the *his*-ePEC structure was determined at relatively low resolution (5.5 Å) from a limited number of particles (7). Thus, we sought a more complete, higher-resolution structure of *his*-ePEC for comparison to *con*-ePEC. Since formation by direct reconstitution gave pause kinetics indistinguishable from actively formed *his*-ePEC (Fig. 1E), we reconstituted the *his*-ePEC on the same scaffold used previously for the hairpin-stabilized *his*-PEC but using an RNA lacking the pause hairpin (Fig. 1B). The resulting *his*-ePECs determined from ~900K polished particles yielded five distinct conformations (Fig. 3A, S4; Table S3). The dominant *his*-ePEC states (73% of the *his*-ePEC particles) were half-translocated with an unfolded-TL but differed from the previously determined *his*-ePEC structure and from each other by varying degrees of swiveling. The most swiveled *his*-ePEC, *his*-ePEC_ufTL1, was the most populated, representing 49% of the total *his*-ePEC particles (Figs. 3A,B). The less swiveled state, *his*-ePEC_ufTL2, represented 24% of the *his*-ePEC particles. The *his*-ePEC_ufTL2 state is likely populated with many intermediate swiveled states, giving rise to the poor resolution of the reconstruction (5.5 Å nominal resolution despite being populated with ~211K particles; Table S3). Thus, we propose that the *his*-ePEC_ufTL states comprise a favored, relatively homogeneous swiveled state (*his*-ePEC_ufTL1) and a population of particles sampling less swiveled states.

**FIGURE 3.**
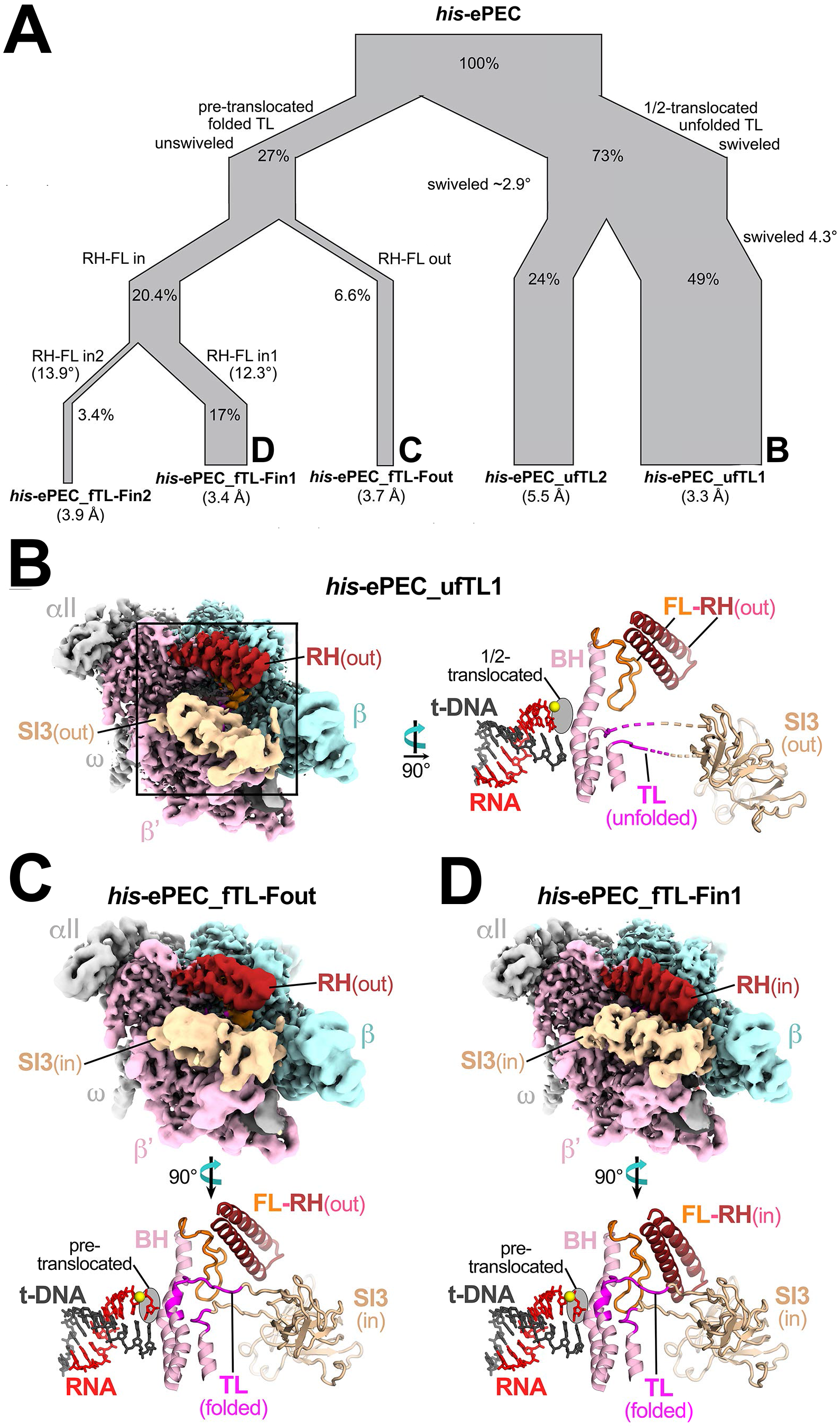
Cryo-EM analysis of directly reconstituted *his*-ePEC. (**A**) Particle-sorting dendrogram of *his*-ePEC conformations. The relative amounts of different *his*-ePEC states and their relationships during particle sorting are represented by the widths and junctions, respectively, of the dendrogram roots. The complete particle sorting analysis for *his*-ePEC is shown in Fig. S5. (**B**) The overall cryo-EM structure and active-site conformation of *his*-ePEC_ufTL1. RNAP subunits and features in the cryo-EM map and rotated active site view are depicted as described for *con*-ePEC_−1_ in Fig. 2B. The half-translocated state of *his*-ePEC_ufTL1 is evident from translocation of the RNA 3’ nt but not its t-DNA partner. The swiveled conformation of *his*-ePEC-ufTL1 is depicted in Fig. 4. (**C**) The overall cryo-EM structure and active-site conformation of *his*-ePEC_fTL-Fout. Graphic details are as described for *con*-ePEC_−1_ in Fig. 2B. (**D**) The overall cryo-EM structure and active-site conformation of *his*-ePEC_fTL-Fin1. Graphic details are as described for *con*-ePEC_−1_ in Fig. 2B.

The remaining ~27% of the *his*-ePEC particles were pre-translocated with the TL folded and SI3 in the closed conformation similar to *con*-ePEC_fTL (Fig. 2C). Three distinct conformations of the RH–FL module could be resolved among these particles. In *his*-ePEC fTL-Fout (6.6% of particles; Fig. 3C), the RH-FL module position was similar to that observed in ECs with an unfolded TL (such as *con*-ePEC_−1_; Fig. 2A) as well as many other *E. coli* RNAP structures. In *his*-ePEC_fTL-Fin1 (17% of *his*-ePEC particles; Fig. 3D), the RH-FL module rotates 12.3° onto the TL (Fig. 3B). In *his*-ePEC_fTL-Fin2 (3.4% of the *his*-ePEC particles), the RH-FL module rotates an additional 1.6° towards the TL. The movement of the RH-FL module in *his*-ePEC_fTL-Fin structures resembles that in a previously reported CTP-bound EC in which the RH-FL module was rotated down over the folded TL (29).

Other than the RH-FL module changes, the folded-TL *his*-ePEC structures were nearly identical (maximum rmsd of 0.577 Å over 3,104 a-carbon positions), giving us an opportunity to assess the effects of the RH-FL conformational changes on contacts between important RNAP structural modules. Rotation of the RH–FL module onto the TL in the *his*-ePEC_fTL-Fin structures generated substantial interface areas between the RH and TL (RH–TL interface area ~105 Å^2^) and the FL–SI3 (~86 Å^2^), whereas there were no contacts (0 Å^2^ interface area) in *his*-ePEC_fTL-Fout (Fig. 4A,B). The RH–SI3 and FL–TL contacts also increased substantially upon RH–FL closing onto the folded TL (Fig. 4B).

**FIGURE 4.**
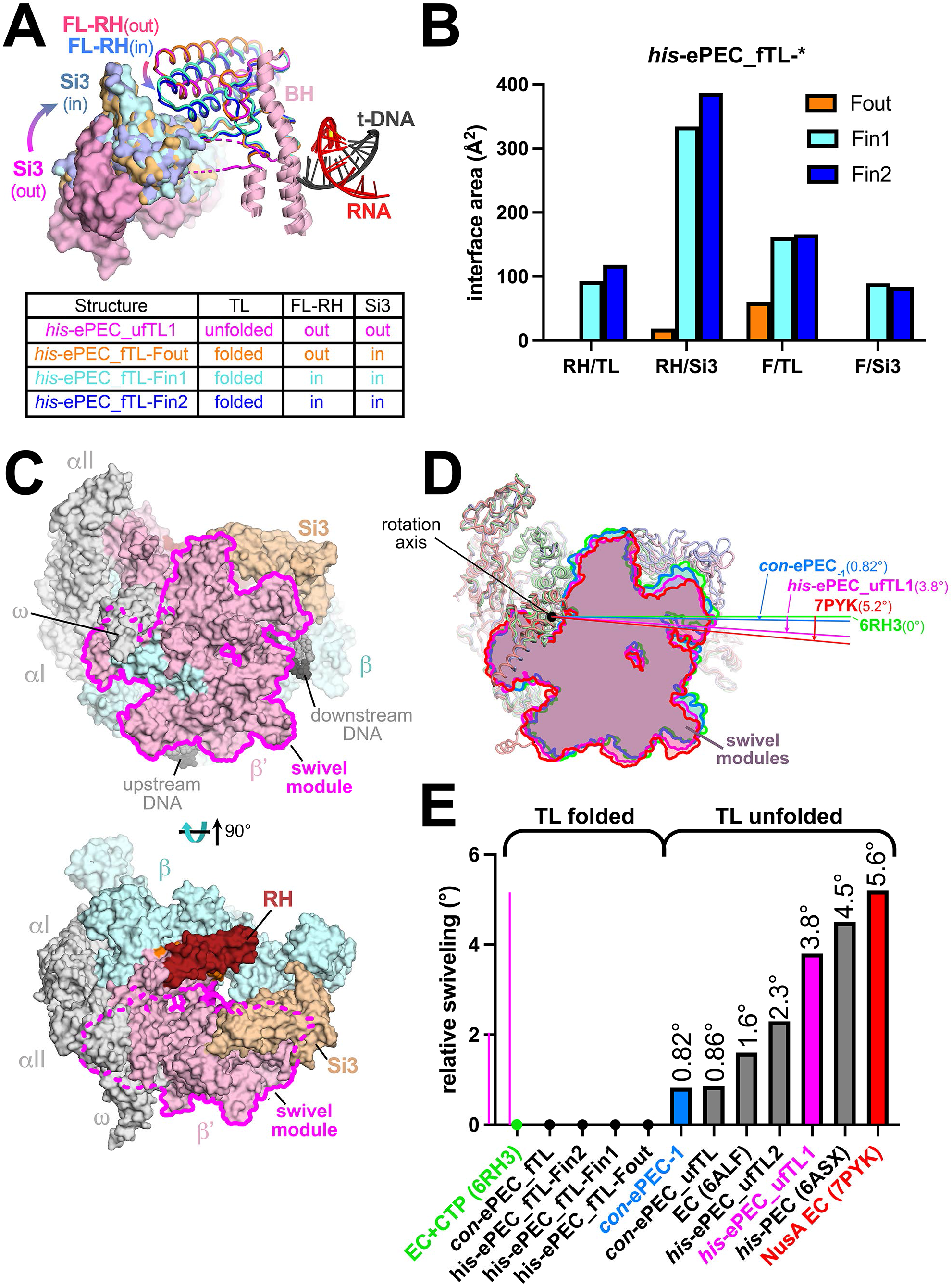
Conformational changes among *his*-ePEC states. (**A**) Comparison of the locations of SI3, RH, and FL in four *his*-ePEC states. *his*-ePEC states are colored as indicated in the figure. The table indicates the status of the TL, the FL–RH modules, and SI3 in each state. (B) The area of the solvent inaccessible interface between RNAP modules in three *his*-ePEC folded-TL states. The interface areas (58) in Å^2^ between two modules separated by a forward slash are plotted for *his*-ePEC_Fout (orange), *his*-ePEC_Fin1 (cyan), and *his*-ePEC_Fin2 (blue). (**C**) The boundary of the swivel module (magenta) comprised of the RNAP shelf, clamp, jaw, β’ C-terminal region, and SI3 when viewed from the ω side (top, corresponding to axis of swivel module rotation) or secondary-channel side (bottom) of RNAP. RNAP subunits are colored as described for con-ePEC_−1_ in Fig. 2B. (**D**) Rotation of the swivel module around the swivel axis for representative ECs and PECs colored as depicted in the figure. (**E**) The angle of swivel-module rotation for ECs and PECs relative to an NTP-bound EC (PDB 6RH3) (30). The differences in rotational angle among the ECs and PECs plotted at 0° are not reliably distinguishable.

The *his*-ePEC_fTL-Fout and *his*-ePEC_fTL-Fin structures all have a folded TL, revealing that FL movement and TL unfolding are not tightly coupled. Retention of the folded TL in the *his*-ePEC_fTL-Fout conformation indicates that RNA–DNA-sequence-determined interactions in the ePEC may inhibit TL unfolding. These interactions may disfavor forward translocation and favor retention of folded TL–nucleic acid interactions in the ePEC as one way for pause sequences to prolong pausing.

### TL unfolding permits RNA–DNA sequence–dependent RNAP swiveling

The ePEC structures described here represent a large range of potential swivel angles (Fig. 4C shows the swivel module in the context of the *con*-ePEC_−1_ structure). To compare swivel angles, we performed structural superpositions based on the RNAP core module (Table S1). As posited by Zhu et al. (30), a structure bound to the incoming CTP substrate with folded TL poised for catalysis (PDB 6RH3) is likely most representative of the catalytically active state and was used as a reference to compare other structures (0° swiveling). Swivel angles computed to be < 0.5° were set to 0° (such small rotation angles are not meaningful because the rotation axis cannot be determined reliably). By contrast, an *E. coli* EC bound to NusA (generally pause-promoting; 7PYK) (30) was found by our analysis to be swiveled 5.6° (Figs. 4D,E). The seven ePEC structures reported here, along with the reference (6RH3) and some other relevant structures (6ALF, *E. coli* EC; 6ASX, *his*-PEC; 7PYK, *E. coli* NusA-EC) (7, 30) exhibit a range of swivel angles from ~0° to 5.6° (Figs. 4D,E).

Previously, we found that swiveling promoted by the *his*-PEC PH (6ASX) allosterically inhibited TL folding by inhibiting SI3 closure (7). Our current results indicate that TL folding in turn inhibits swiveling; all of the compared structures containing a folded TL were unswiveled (~0° swiveling; Fig. 4E). Unfolding of the TL allows swiveling to occur, but to varying degrees (Figs. 4D,E). All of the swiveled *his*-ePEC structures are substantially more swiveled (2.3°–3.8°) than the swiveled *con*-ePEC structure (~0.8°), suggesting that swiveling is RNA–DNA-sequence dependent.

### Cys-triplet reporter verifies large differences in *con*-ePEC and *his*-ePEC states

The strong pre-translocation bias of the *con*-ePEC differs from the half-translocated, swiveled states that dominated the directly reconstituted *his*-ePEC. We next asked if these differences were also evident under conditions of active transcription in solution, using longer, fully complementary scaffolds that lack perturbing influences of an artificial transcription bubble and short downstream DNA present in cryo-EM complexes (Figs. 5A and S5). We first used a Cys-triplet reporter (CTR) assay to distinguish closed and swiveled positions of SI3 in *con*-ePEC and *his*-ePEC (Figs. 5B and S5B) (34). In the CTR assay, the closed SI3 forms a disulfide between engineered β’1051C in the SI3 hairpin loop and β’671C at the tip of the RH. The swiveled SI3 instead forms a β’1051C disulfide with β267C in SI1 and the two disulfides are distinguishable by non-reducing SDS-PAGE (Fig. S5C). The ratio of the closed to swiveled disulfide, denoted SI3 positional bias (SPB), is a relative measure of influence of transcription complex conformation on SI3 position (34).

**Figure 5.**
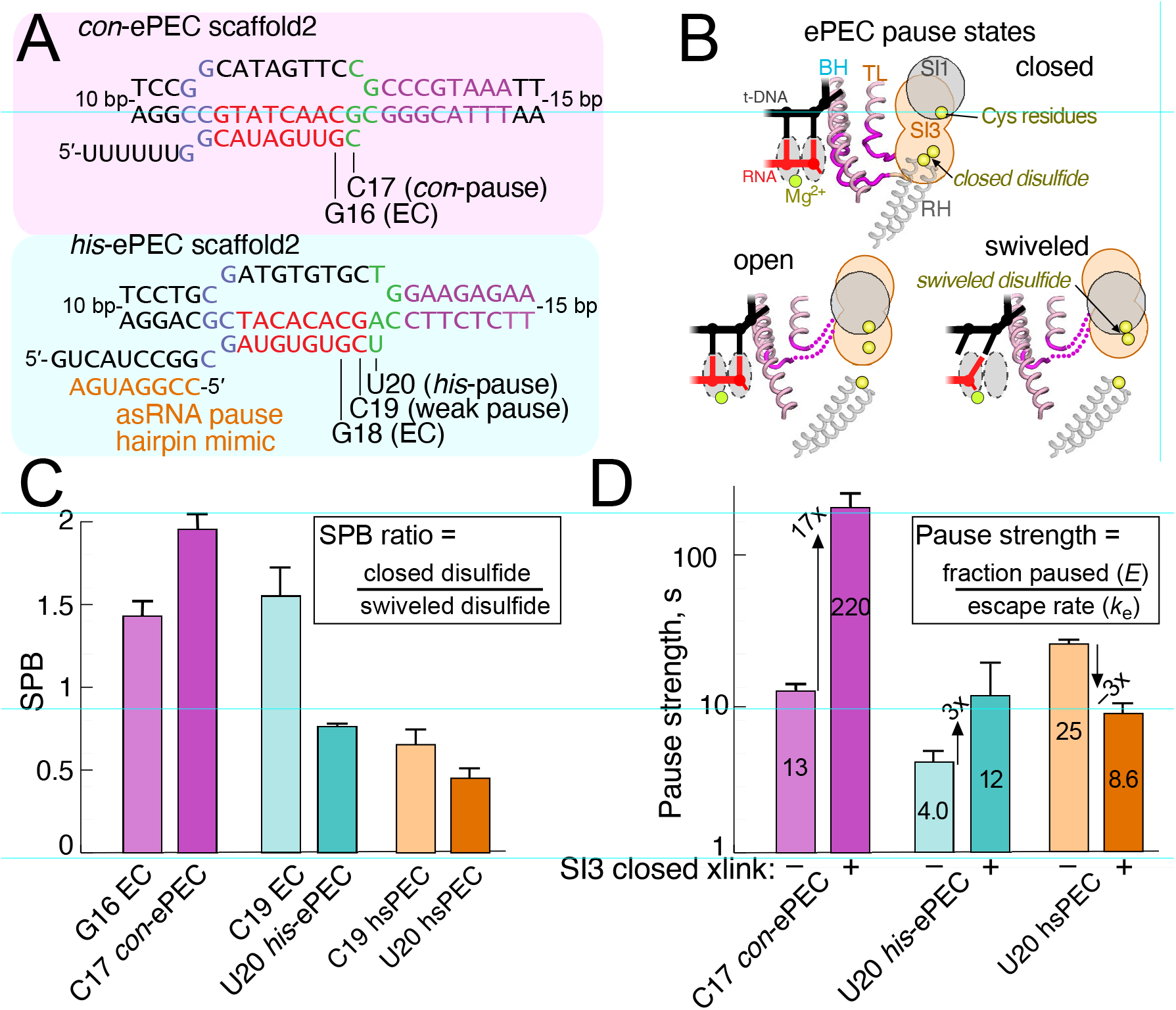
Measurement of SI3 location by Cys-triplet reporter (CTR) assay and pause kinetics for *con*-ePEC and *his*-ePEC. (**A**) The scaffold sequences used for CTR and pausing assays of *con*-ePEC and *his*-ePEC. These scaffolds are fully complementary and contain sufficient duplex downstream DNA to avoid perturbing effects on translocation register (see Fig. S5A for complete sequences). (**B**) Diagrammatic representation of Cys residue locations near the active site of RNAP for CTR assay in SI3 closed, open, and swiveled states (34). (**C**) SI3 positional bias (SPB) as calculated from the CTR assay for ECs and PECs. A higher SPB indicates greater occupancy of the SI3 closed conformation. Note that absolute SPB values do not correspond directly to SI3 closed-to-swiveled ratios but that changes in SPB roughly approximate the changes in these ratios (34). The increase in SPB for *con*-ePEC vs *con*-ePEC_−1_ and decrease in SPB for *his*-ePEC vs. *his*-ePEC_−1_ indicates that *con*-ePEC favors the closed SI3, presumably pre-translocated, conformation related to an EC whereas his-ePEC favors the swiveled SI3, presumably half-translocated conformation, related to an EC. (D)). Pause strengths of paused ECs with and without a disulfide crosslink that restrains SI3 in the closed position (closed SI3 xlink). The *his*-ePEC results depicted here are from experiments reported previously (34).

To ask if SPB differed in *con*-ePEC and *his*-ePEC, we reconstituted the two ePECs by active elongation from an EC positioned one nucleotide upstream of the pause site (G16 for *con*-ePEC or C19 for *his*-ePEC). *his*-ePEC_−1_ is known to also pause weakly (35). When compared to the −1 ECs, SPB increased for *con*-ePEC and decreased for *his*-ePEC (Fig. 5C). Consistent with previous results, formation of a PH mimic using an antisense RNA further decreased SPB for *his*-ePEC. These results indicate that SI3 became more biased toward the closed position in *con*-ePEC, consistent with the pre-translocated *con*-ePECs observed by cryo-EM. Conversely, SI3 became more swiveled in the *his*-ePEC, also consistent with predominant occupancy of swiveled states observed by cryo-EM. Using an assay orthogonal to cryo-EM and scaffolds devoid of potentially perturbing influences, these results confirm that active *con*-ePEC and *his*-ePEC formed on scaffolds occupy predominantly pre-translocated and predominantly swiveled conformations, respectively.

The disulfides used in the CTR assay also can be used individually to bias ECs toward the SI3-closed or swiveled conformations (34). To ask if biasing SI3 toward the closed conformation would affect the pause strength of *con*-ePEC and *his*-ePEC similarly or differently, we measured the effect of the closed disulfide on pause kinetics (Figs. 5D and S5D,E). Trapping the closed SI3 with a disulfide dramatically enhanced the pause strength and lifetime of the *con*-ePEC (Fig. 5D and S5D). In contrast, the same disulfide modestly increased the strength of the *his*-ePEC and decreased the strength of the PH-stabilized *his*PEC (Fig. 5D and S5E). These contrasting effects of the closed SI3 disulfide are consistent with bias toward the pre-translocated state in the *con*-ePEC that is greatly strengthened by the disulfide and a bias toward the swiveled conformation in the *his*-ePEC that is countered by the disulfide, resulting in a modest effect on pause lifetime.

### A translocation register assay confirms different *con*-ePEC and *his*-ePEC states

As a further test of differences in translocation bias of the *con*-ePEC and *his*-ePEC, we next examined reconstituted ePECs using an assay that traps ECs or PECs in their current translocation register with the phage HK022 Nun protein (27, 36). This assay reports the post-translocated state based on its capacity for nucleotide addition and the pre-translocated state based on its sensitivity to pyrophosphorolysis (Fig. 6A; S6). For this assay, we reconstituted *con*-ePEC and *his*-PEC on scaffolds similar to those used for cryo-EM but fully complementary to eliminate possible perturbation of translocation register caused by non-complementarity (Fig. S6A). It was unclear how the half-translocated PEC would be affected by Nun. We formed ePEC_−1_ by reconstitution, bound to Ni^2+^-NTA beads via an RNAP His_10_ tag, extended to ePECs by incubation with CTP (for *con*-ePEC) or UTP (for *his*-ePEC), treated with Nun, washed away NTPs, and then incubated with 1 mM GTP or 2.5 mM PPi to assay translocation register (Fig. S6B). Nun-treated *con*-ePEC was more sensitive to pyrophosphorolysis and incorporated less GTP, whereas the opposite was true for *his*-ePEC and to an even greater extent for the hairpin-stabilized *his*PEC (Fig. 6B and S6D–G). Since the *his*PEC is known to be half-translocated, we infer the Nun-bound, half-translocated EC can react with NTP substrate. Taken together, these data are thus fully consistent with the cryo-EM structures suggesting that *con*-ePEC is principally pre-translocated and *his*-ePEC is principally half-translocated.

**FIGURE 6.**
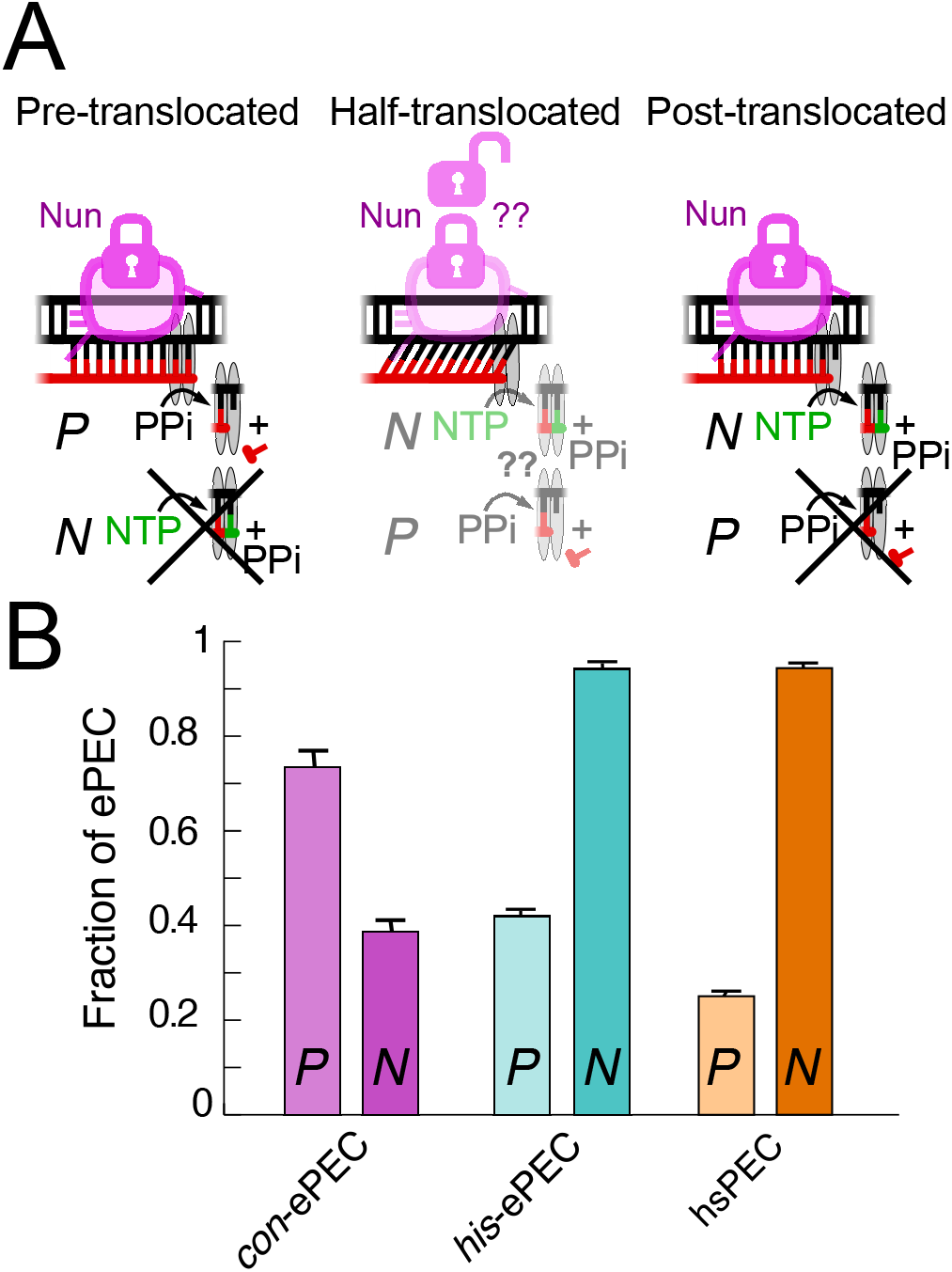
Translocation registers of *con*-ePEC and *his*-ePEC determined by Nun-locked assay (36). (A) Diagrammatic representation of Nun-locked translocation assay for pre-, half-, and post-translocated states. Nun locks translocation register via interactions with the upstream fork junction and downstream duplex DNA. Pre-translocated ECs are susceptible to pyrophosphorolysis but not nucleotide addition. Post-translocated ECs are susceptible to nucleotide addition but not pyrophosphorolysis. Whether Nun locks the half-translocated state and its subsequent reactivity are uncertain, but results presented here are consistent with either reactivity with NTP not PPi, or Nun-induced conversion to a post-translocated state reactive with NTP but not PPi. (**B**) Fractions of PECs reactive with PPi (P) or NTP (N) after incubation with Nun. Greater reactivity with PPi and lesser reactivity with NTP of *con*-ePEC that *his*-ePEC or hairpin stabilized (hs) *his*-hsPEC is consistent with greater occupancy of pre-translocated register for the *con*-ePEC and greater occupancy of half-translocated registers by *his*-ePEC and *his*-hsPEC as also observed by cryo-EM analyses. Data are average and s.d. of 3 independent replicates from gels shown in Fig. S6C–E.

### Multiple ePEC states can explain biphasic pause kinetics and RNA–DNA sequence effects on elemental pausing

The identification of multiple co-existing ePEC states provides a potential explanation for multiphasic ePEC pause kinetics that were found previously not to involve backtracking (19). Sequence-dependent differences in the preferred states also may help explain how multipartite RNA–DNA sequences modulate pausing. To explore these explanations, we considered how the different ePEC states may be connected structurally and energetically (Figs. 7A, S7, S8, and S9). The pre-translocated–folded TL (closed SI3) state should be the first to form when the EC arrives at a pause site because it is directly generated by the nucleotide-addition reaction. We thus assigned this state as the first paused state, most likely the fTL–Fin2 state with the tightest F-loop–TL interaction (state A, Fig. 7A). Pause entry directly into the predominant *his*-ePEC fTL–Fin1 state (state B) also is possible.

**Figure 7.**
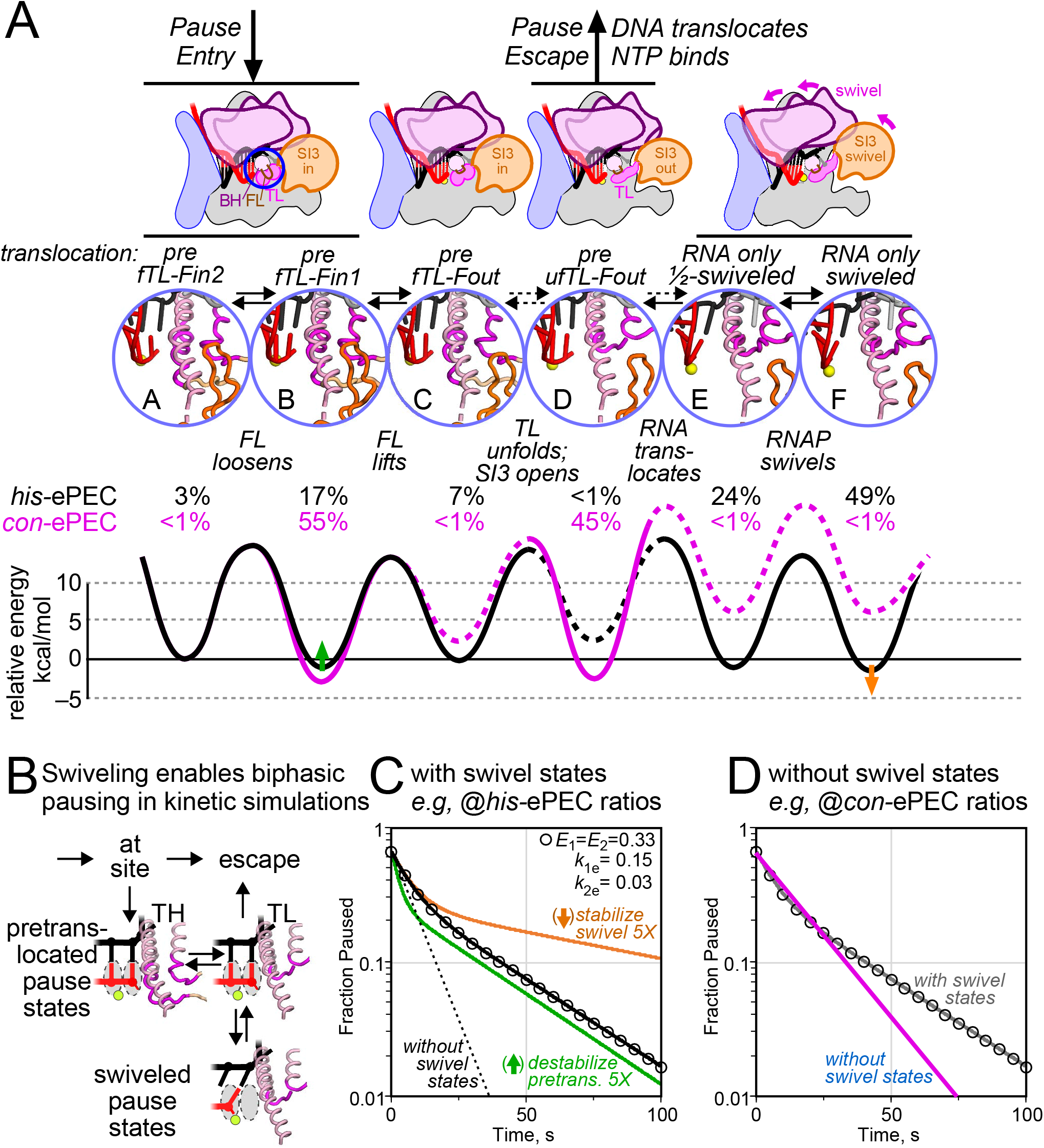
The multiple intermediate model of elemental transcriptional pausing. (**A**) Distinct ePEC intermediates revealed by *con*-ePEC (intermediates B and D) and *his*-ePEC (intermediates A, B, C, E, and F) cryo-EM and their deduced energetic relationships (see Table 1 for comparison to *con*-ePEC and *his*-ePEC intermediate names). The relative stabilities of the intermediates were calculated from cryo-EM particle distributions (percentages shown in black or magenta) assuming a constant rapid forward rate (100 s^-1^, left to right) and fitting reverse rates to observed intermediate occupancies using Kintek Explorer (59) (Methods). The relative energy levels but not the absolute rates are constrained by the distributions of ePEC states. (**B**) The swivel model of pausing accounts for biphasic pause escape kinetics. (**C**) Fitting of the swivel model to a typical set of biphasic pause escape kinetics. Fraction paused RNA remaining as a function of time was calculated using the parameters listed and a two-exponential rate of decay (open circles). A kinetic model that included off-pathway swiveled states E and F and fixed observed ratios of states A–F for *his*-ePEC was fit to these data successfully (black line) using Berkeley Madonna (60) whereas a kinetic model lacking off-pathway swiveled states was unable to fit these data (dotted line) (Methods). Five-fold changes in the stabilities of pretranslocated state B (green line, corresponding to green arrow in panel A) or swiveled state F (orange line, corresponding to orange arrow in panel A) altered either the slow or slower components of the biphasic pause escape rates. Kinetic constants used to generate these data are shown in Fig. S10. (**D**) Fitting of a kinetic model lacking off-pathway swiveled states (fixed observed ratios of states A–F for con-ePEC was unable to give biphasic kinetics (magenta line; predicted data are same as for panel C). Fitting that allowed occupancy of swiveled states was readily able to generate a good biphasic fit (gray line). Kinetic constants used to generate these data are shown in Fig. S10.

The active site would then relax by movement of the FL and RH away from the folded TL (state C). This movement of the FL and RH would allow subsequent unfolding of the TL and shifting of SI3 to the open position to yield a pre-translocated complex ready for translocation (state D). We postulate that full translocation of both RNA and DNA from state D would create a post-translocated EC ready for nucleotide addition and would constitute pause escape. However, translocation of only the RNA and not the DNA would form a half-translocated paused state associated with swiveling (state E). It is unclear if swiveling leads to half-translocation, or vice versa, or if swiveling and half-translocation are mutually reinforcing. Further swiveling would yield the swiveled ePEC observed in the *his*-ePEC that can be stabilized by pause RNA hairpin formation (state F).

Because we knew the equilibrium distributions of the different states for the *con*-ePEC and *his*-ePEC from the cryo-EM particle distributions, we could estimate the relative energetics of their interconversion. The *con*-ePEC principally occupies states B and D (magenta energy diagram; dotted lines indicate unoccupied states for which only limits on the energy diagram can be ascertained). The height of energy barriers between states dictates their rates of interconversion, whereas their relative occupancies at equilibrium depend on the relative positions of the troughs. The *his*-ePEC, in contrast, occupies states A, B, C, E, and F (black energy diagram).

This analysis provides a parsimonious hypothesis that can explain biphasic pause escape kinetics by formation of swiveled states (Fig. 7B). To illustrate this point, we fit a kinetic model with fixed ratios of the paused states to a calculated dataset of biphasic pause kinetics (black circles, Fig. 7C). Fitting was accomplished using predictions of numerically integrated rate equations and least-squares minimization (Methods and Fig. S9). The *his*-ePEC state ratios, which include the swiveled states, readily fit the biphasic pause dataset (black line, Fig. 7C). However, a *his*-ePEC kinetic model lacking swiveled states was unable to generate biphasic pausing behavior (dotted line, Fig. 7C). The same was true for a *con*-ePEC kinetic model lacking swiveled states (blue line, Fig. 7D). Addition of swiveled states to the *con*-ePEC model readily allowed generation of a good fit to the biphasic pause dataset (gray line, Fig. 7D). Biphasic kinetics are possible with the swiveled states because they provide an additional level of off-pathway states that ePECs can enter and escape from more slowly. In this model, ePECs entering these swiveled states must unswivel to be able to complete translocation. Kinetic models lacking this additional level of off-pathway states predict a single rate of pause escape.

Small changes (by factors of 5) in the stabilities of the pause states predicted obvious changes on distinct phases of pause escape (green and orange lines, Fig. 7C). These changes illustrate how small changes caused by different RNA–DNA sequences or interactions with transcription factors may modulate pause kinetics.

We emphasize that the off-pathway effect of the swiveled states is a hypothesis. It is attractive because it readily explains what has been otherwise mysterious: how biphasic pause escape kinetics arise without involvement of backtracking. However, other explanations are possible. For example, the fTL-Fin2 state (state A, Fig. 7A) could be off-pathway and changes in its rates of formation and collapse could generate biphasic kinetics. We favor the swivel hypothesis because it is parsimonious. Swiveling is only observed when the TL is unfolded (Fig. 4E). If nucleotide addition cannot be completed once RNAP swivels, then swiveling conveniently enables modulation of pause strength by interactions that stabilize the swiveled state(s). Such stabilization is known to occur by formation of a PH in the RNA exit channel of RNAP or by binding of NusA (7, 9, 30).

## DISCUSSION

RNA–DNA sequence–dependent pausing underpins transcriptional regulation. Most investigators now agree that RNAP responds to pause signals by initially entering an offline, elemental paused state that is neither backtracked nor stabilized by nascent RNA structures or diffusible regulators. Half-translocation (RNA but not DNA) has been associated with elemental pausing and swiveling with pause stabilization (7, 9, 28, 30), but a definitive structural description of the ePEC has remained elusive. By comparing two ePECs formed on different nucleic acid sequences, we found that the ePEC actually comprises a family of distinct paused states. Different ePEC conformations dominate for different pause signals. A strong consensus pause signal was predominantly pre-translocated whereas a weaker signal found in the *his* biosynthetic operon leader region was predominantly swiveled even in the absence of the swivel-stabilizing, nascent PH. However, for each ePEC multiple distinct conformations could be resolved by cryo-EM. This view of the ePEC as a family of interconverting states, rather than a single species, resolves some mysteries about elemental pausing, has implications for both transcriptional regulation and our understanding of the fundamental mechanism of nucleotide addition by RNAP, and raises new questions for future study.

### Multiple ePEC states increase regulatory options

The ability of ePECs to occupy multiple conformations creates opportunities for differential regulation of transcriptional pausing. Both RNA–DNA sequences and diffusible regulators may modulate the dwell time of RNAP at a pause as well as the fraction of RNAP molecules that isomerize into the ePEC versus continuing rapid transcription. Together the pause fraction and pause dwell time determine pause strength, which is the mathematical product of the two parameters (37, 38). When a single pause state exists, pause fraction and dwell time are interdependent. This relationship has been observed for short-lived pauses by single-molecule methods at 23 °C (39). The existence of multiple interconverting pause states, on the other hand, could allow unlinked modulation of pause fraction and pause dwell time, creating increased regulatory flexibility.

For example, transcriptional regulators may modulate the dwell time of pauses by stabilizing or destabilizing swiveling without changing the fraction of RNAP molecules that pause. This is precisely how PHs are thought to operate. Formation of the ePEC allows time for nascent PH formation. PH–RNAP interactions may then stabilize the swiveled state, which increases pause dwell time to allow time for regulator interactions. In *E. coli*, ribosomes may then initiate translation on the nascent RNA and subsequently melt the PH, destabilizing the swiveled state and triggering pause escape. These steps synchronize translation to RNAP movement over a terminator thus linking translational status to transcriptional attenuation (40).

Other cellular components that interact with nascent RNA may substitute for the ribosome in this type of indirect regulation of pausing (i.e., via RNA structure). For example, small molecules can bind the nascent RNA during riboswitch attenuation (41). The unlinking of pause dwell times from pause fraction that is possible due to the existence of multiple ePEC states allows these mechanisms to operate with high efficiency by ensuring that most RNAPs enter pauses that can be modulated by different interactions. Direct modulation of the swiveled state via contacts to RNAP also is possible. For example, modulation of pausing by the universal transcription factor NusG and its paralogs like RfaH may operate via enhancement or suppression of swiveling (14).

Conversely, other interactions with RNAP may trap the ePEC in a non-swiveled but nonetheless significant pause. For example, interactions of the RNA–DNA hybrid with RNAP may stabilize the pre-translocated paused state so much that readthrough of the pause becomes hard to detect (4, 19, 42). The opportunity to stabilize non-swiveled pauses afforded by multiple ePEC states may be especially important for halting antiterminated ECs whose swiveling is suppressed. In antiterminated ECs, RNAPs are modified by diverse regulators that promote transcript elongation, including by inhibiting swiveling (43–46). Antiterminated ECs still must terminate at the ends of functional transcription units. This termination appears to be accomplished by specialized intrinsic terminators. Pausing is the first step in intrinsic termination. It is possible that pausing in the pre-translocated register may be crucial for function of these specialized terminators where swiveling would remain suppressed by the antitermination modifications.

### Regulation of multiple pause states may be analogous to regulation of multistep transcription initiation

Decades of work have elucidated how RNAP initiates transcription at multipartite promoters (e.g., containing UP, −35, −10 elements, etc.) by a multistep process involving sequential intermediate states from promoter recognition to promoter escape (47, 48). Each step is subject to differential regulation whereby different transcription factors and different promoter sequences may alter the overall rate of initiation by affecting different steps in the multistep initiation process (47, 49). For example, some factors or sequences may increase initial promoter melting and others may stabilize the open complex.

The multiple paused–state model (Fig. 7) suggests regulatory opportunities that are remarkably parallel to the more fully understood mechanism and regulation of transcription initiation. Like the multipartite promoter, pause sequences also are multipartite (e.g., upstream RNA structures and sequences of the upstream fork junction, RNA–DNA hybrid, downstream fork junction, and downstream DNA duplex; Fig. 1B) (6, 19). Different regulators affect pausing through different interactions with these sequences (e.g., NusA with RNA structures, NusG with upstream fork junction and nt-DNA) (8, 9, 50). The multiple ePEC states defined here thus suggest regulatory opportunities for transcriptional pausing that parallel the well-defined regulation of multistep transcriptional initiation. Unlike for initiation, the roles of different parts of the multipartite pause signals in controlling steps in the pause mechanism are unknown (*e.g*., it is unclear which sequence elements control swiveling). Elucidating these connections will be an important focus for future research.

### Swiveling and half-translocation: steps in the NAC or off-pathway paused states?

Whether swiveling and half-translocation could be sub-steps in the ordered events that occur during every NAC or only happen in offline paused states remain open questions. Swiveling has been observed in subpopulations of post-translocated ECs formed on nominally non-pausing scaffolds by cryo-EM (30). It also has been proposed to aid normal translocation of the RNA–DNA hybrid by ratchet-like action, at least in some bacterial RNAPs (51). Half-translocation has been detected in time-resolved crystallography of nucleotide addition by viral RNA-dependent RNAPs and proposed to be an intermediate in the NAC (52).

Our detection of these states in equilibrium with pre-translocated ePECs does not directly address whether half-translocation or swiveling can occur during the non-paused NAC of multisubunit DNA-dependent RNAPs. However, the changes in RNAP substructures around the active site that are observed in the swiveled state, including a slight bending of the BH (7), the loosened locations of the RH and FL (Fig. 3B), and changes in the clamp location, appear incompatible with their conformations in the NTP-bound, TL-folded state poised for nucleotide addition. Thus, a parsimonious interpretation is that swiveling occurs in offline ePECs and neither is a necessary aid to translocation nor compatible with rapid catalysis during the NAC (30, 53). The observation of swiveled subpopulations in ECs formed on non-pause-site scaffolds (30) could reflect slow formation of ePEC conformations in halted ECs not relevant on the time scale of active transcription or rapid, transient sampling of swiveled states in unfolded-TL ECs that only become long-lived in PECs.

Whether half-translocation occurs as an intermediate in every round of the NAC may not be easily resolved since the half-translocated state could be extremely short-lived. It also may not be mechanistically important. Inhibition of DNA translocation after RNA translocation in the half-translocated state clearly generates a rate-limiting barrier to escape from the ePEC family of states. This interpretation is directly validated by the finding that nt-strand substitutions or RNAP mutants that weaken capture of the translocated +1 non-template base in the so-called CRE pocket can greatly alter pause dwell times (22, 23, 54). However, the rate of transcript elongation over long DNA segments is not significantly affected by substitutions in the CRE pocket that affect pausing (54). Molecular dynamics simulations of translocation do not reveal significant occupancy of the half-translocated state (55, 56), although further application of this approach could give additional insight. Normal thermal fluctuations of the EC could allow either RNA-first or concerted RNA–DNA translocation to occur, making the half-translocated state a possible but inconsequential intermediate in the normal NAC.

Finally, we note that our attempt at manual time-resolved detection of paused states did not yield clear evidence of intermediates that precede the apparently equilibrated states formed by direct reconstitution. For example, we did not detect complexes retaining pyrophosphate. Direct reconstitution of the *his*-ePEC revealed states similar to those formed by freeze-capture of *con*-ePECs after nucleotide addition. Nonetheless, the approach of capturing intermediates shortly after nucleotide addition holds promise and may provide insight into how ePECs form or how ECs bypass pauses if new methods able to achieve sub-second time resolution are applied (57).

## ACKNOWLEDGEMENTS

We thank members of the Darst-Campbell and Landick Laboratories for experimental advice and helpful discussions, and M. Ebrahim, J. Sotiris, and H. Ng at The Rockefeller University Evelyn Gruss Lipper Cryo-electron Microscopy Resource Center for help with cryo-EM data collection. This work was supported by NIH grants R35 GM118130 to S.A.D and R01 GM38330 to R.L.

## Notes

### Competing Interest Statement

The authors have declared no competing interest.

